# Plotgardener: Cultivating precise multi-panel figures in R

**DOI:** 10.1101/2021.09.08.459338

**Authors:** Nicole E Kramer, Eric S Davis, Craig D Wenger, Erika M Deoudes, Sarah M Parker, Michael I Love, Douglas H Phanstiel

## Abstract

**T**he R programming language is one of the most widely used programming languages for transforming raw genomic data sets into meaningful biological conclusions through analysis and visualization, which has been largely facilitated by infrastructure and tools developed by the Bioconductor project. However, existing plotting packages rely on relative positioning and sizing of plots, which is often sufficient for exploratory analysis but is poorly suited for the creation of publication-quality multi-panel images inherent to scientific manuscript preparation. We present plotgardener, a coordinate-based genomic data visualization package that offers a new paradigm for multi-plot figure generation in R. Plotgardener allows precise, programmatic control over the placement, aesthetics, and arrangements of plots while maximizing user experience through fast and memory-efficient data access, support for a wide variety of data and file types, and tight integration with the Bioconductor environment. Plotgardener also allows precise placement and sizing of ggplot2 plots, making it an invaluable tool for R users and data scientists from virtually any discipline.

**Availability:** Package: https://bioconductor.org/packages/plotgardener

Code: https://github.com/PhanstielLab/plotgardener

Documentation: https://phanstiellab.github.io/plotgardener/

## Rationale

The increasing size, complexity, and sheer volume of multi-omic data sets has created a dire need for tools to efficiently visualize, interpret, and communicate the underlying biological signals present in these data. Towards this end, genome browsers, including the UCSC Genome Browser and IGV, have revolutionized our ability to investigate genomic data in a rapid and intuitive fashion, using a stacked linear representation of a wide variety of data types and annotations^1–8^. Recently, more specialized browsers like Juicebox^9^ and HiGlass^10^ have increased the ability to visualize non-linear data types, such as 3D chromatin contact frequency^11,12^. Furthermore, an ever-increasing array of programmatic libraries and browser APIs now allow code-based, integrated data analysis and construction of browser tracks, which has improved reproducibility and automation^13–17^.

While these tools have been transformative for data exploration, they are largely based on single-panel figures and vertical stacking of genomic tracks and are often ill-suited for the generation of complex multi-panel figures that include both genomic and non-genomic plot types. Such complex figures are often critical for evaluating the underlying biology and are almost always used to present multi-omic data in publications. Thus, a tool specifically designed to programmatically create and arrange publication-quality multi-panel figures is critical to extend the rigor, reproducibility, and clarity of scientific data visualizations.

Currently existing R packages like patchwork^18^, cowplot^19^, gridExtra^20^ and Sushi^21^ (which was developed by our group) can be used to arrange multi-panel plots. However, these layout packages use relative positioning to place plots, giving users little control over precise sizing and arrangement. Figures generated with these tools often need finishing in graphic design software such as Adobe Illustrator^22^, Inkscape^23^, PowerPoint^24^, and Keynote^25^. In addition to the cost of purchasing proprietary graphic design software and the steep learning curve often associated with their use, generating multi-panel figures with these software requires non programmatic, manual user interactions, a labor intensive process that decreases reproducibility.

Here we introduce plotgardener, an R package for absolute coordinate-based plot placement and sizing of complex multi-panel plots. This paradigm gives users precise control over size, placement, typefaces, font sizes, and virtually all plot aesthetics without the need for graphic design software. Plotgardener (1) supports a vast array of genomic data types, (2) allows precise placement and sizing of genomic and non-genomic figures, (3) is tightly integrated with the Bioconductor environment^26^, and (4) is optimized for speed and user-experience. The code is open source, extensively documented, and freely available via GitHub and Bioconductor.

## Philosophy

The defining feature of plotgardener that separates it from virtually all other genomic visualization tools is that it allows exact sizing and placement of plots using an absolute, coordinate-based plotting system **(Fig. 1)**. Each plot, axis, and annotation is placed independently according to user-specified positions and dimensions. Each plot or feature extends from edge to edge of the defined coordinates, allowing for precise control and perfect alignment of plots. Rulers and guidelines can be temporarily added for ease of plotting and then removed prior to file generation. Adding additional plots does not shift or resize existing ones, so figures can be built incrementally and adjusted without affecting other figure panels, allowing rapid and easy construction of publication-quality multi-panel figures.

**Fig. 1.**
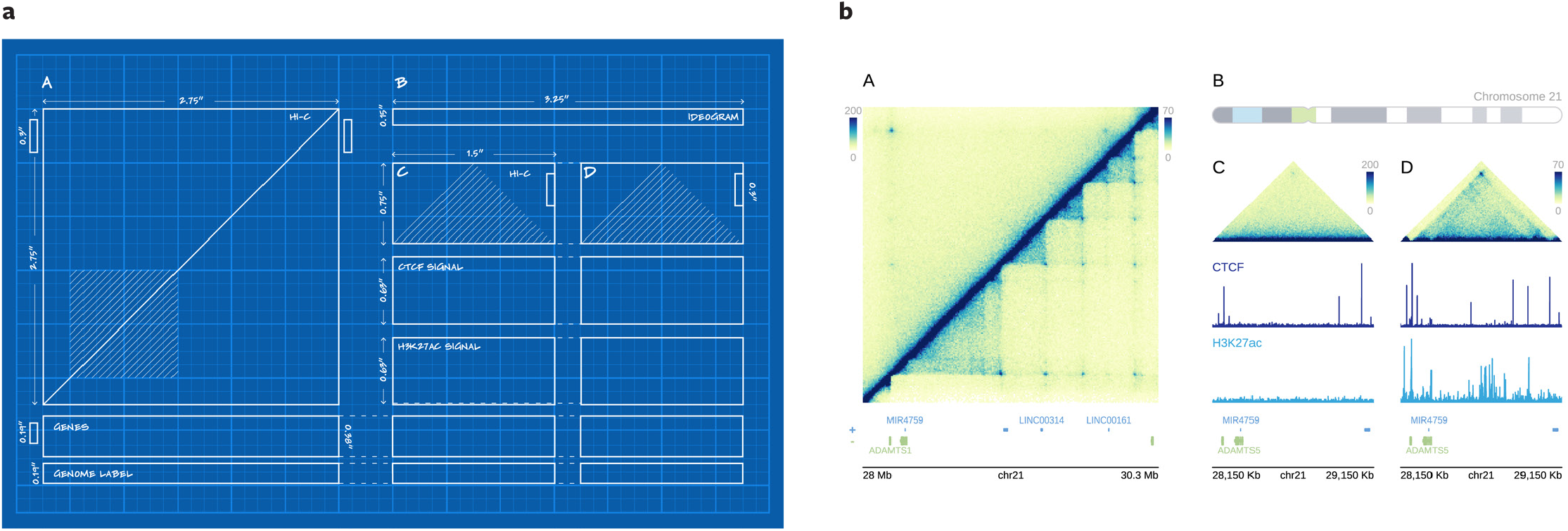
Plotgardener uses a coordinate-based plotting system to size and arrange plots. **a**, Blueprint outline of a multi-omic figure to be created with specified dimensions and placements on a defined page. **b**, Multipanel, multi-omic figure programmatically created with plotgardener using the sizing and placement coordinates from (a). The plotgardener functions used to create this figure include pageCreate, plotHicSquare, annoHeatmapLegend, plotGenes, annoGenomeLabel, plotIdeogram, plotHicTriangle, plotSignal, and plotText. Code to reproduce this plot is included in the plotgardener package.

## Data types

Plotgardener can display a vast array of genomic data types which can be provided as either external files or R data classes. Plotgardener has 45 functions for plotting and annotating diverse genomic data types, including genome sequences, gene/transcript annotations, chromosome ideograms, signal tracks, GWAS Manhattan plots, genomic ranges (e.g. peaks, reads, contact domains, etc), paired ranges (e.g. paired-end reads, chromatin loops, structural rearrangements, QTLs, etc), and 3D chromatin contact frequencies. Plotgardener automatically recognizes and reads compressed, indexed file types including “.bam”, “.bigwig”, and “.hic”, allowing for rapid and memory-efficient reading and plotting of large genomic data. **Fig. 2** displays the runtime required to read and plot various types of genomic data. Even with file sizes exceeding 50 GBs, plotgardener can read and plot data in under a second. Multiple classes of R objects are supported, including “data.frame”, “data.table”, “tibble”, “GRanges”, and “GInteractions”. Plotgardener automatically detects whether the input is a file path or an R object and handles them accordingly, providing a seamless and flexible experience for the user.

**Fig. 2.**
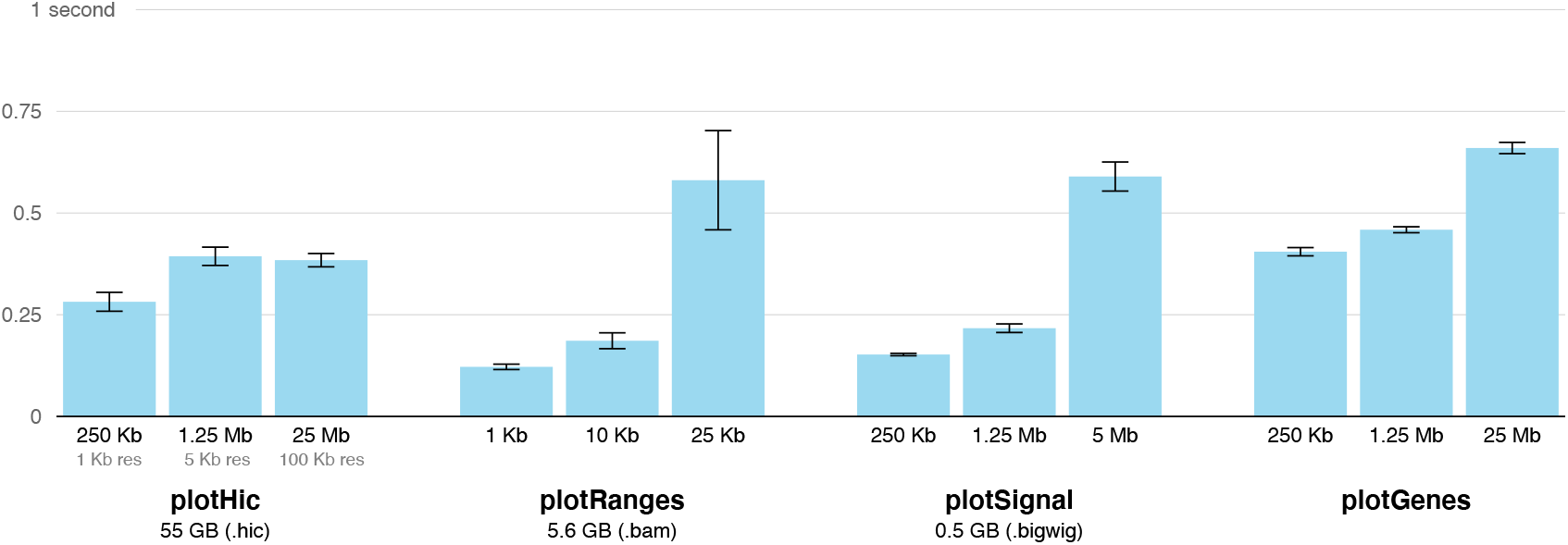
Plotgardener function runtimes. A bar plot depicting mean runtimes for reading and plotting genomic data across various sizes and resolutions using plotHicSquare, plotRanges, plotSignal, and plotGenes functions. Times were calculated for 20 randomly chosen gene regions for each bar. Error bars indicate standard error. File sizes for the input data are indicated below each set of bars for each function: plotHicSquare (55 GB .hic file), plotRanges (5.6 GB .bam file), plotSignal (0.5 GB .bigwig), plotGenes (NA, data stored as an internal object).

## Bioconductor integration

Plotgardener is tightly integrated with the Bioconductor ecosystem^26^, making it compatible with many existing workflows. It has 29 built-in genomes and associated annotations but can easily accommodate custom genomes and annotations using Bioconductor TxDb^27^, OrgDb^28^, and BSgenome^29^ packages and/or objects. Plotgardener leverages these annotation resources on behalf of the user to obtain and plot chromosome sizes, gene and transcript structures, and nucleotide sequences. By preconfiguring the genome builds and associated feature data, plotgardener allows users to focus their attention on layout and to quickly visualize their data rather than spending time and effort on curation and organization of sequences and genome annotations.

## User experience

Plotgardener includes a variety of user-friendly features to maximize ease of use for both novices and experienced R programmers. We describe just some of these features here. Parameters can be set within each function call or passed in a pgParams object for more efficient code. Genomic coordinates can be set either by supplying the chromosome, start, and end position or by providing a gene name (e.g. IL1B), reference genome name (e.g. “hg19”), and optional base pair window around the gene (e.g. 50,000 bp). Resolution of Hi-C contact matrices, signal tracks, and gene tracks are automatically determined based on the genomic range being plotted, but can be overwritten if desired. When genomic regions are too large to label all genes, plotGenes and plotTranscripts will choose which genes/transcripts to label based on frequency of appearance in publications. Users can provide their own priorities or select individual genes to highlight with text and colors. A “colorby” function allows users to flexibly color genomic features by quantitative and qualitative attributes. Plotgardener is open source, version controlled, and extensively documented via articles and vignettes (https://phanstiellab.github.io/plotgardener/).

## ggplot and beyond

In addition to its included functions for plotting and annotating genomic data, plotgardener allows for the absolute sizing and placement of non-genomic plots within a plotgardener page. Users can make multi-panel figures seamlessly by integrating and aligning plotgardener and non-plotgardener plots or create coordinate-based layouts entirely composed of external plot types and objects. For example, plotgardener was used to arrange and add text annotations to the ggplot2 plot objects featured in **Fig. 3**. Plotgardener intuitively sizes, arranges, and overlays plots, text, and geometric objects to make complex figure arrangements beyond basic grid-style or relative layouts. We are actively developing the package and potential future additions include more plotting functions, templates for common arrangements, convenient functions for multiplotting, enhanced ggplot2 integration, and more.

**Fig. 3.**
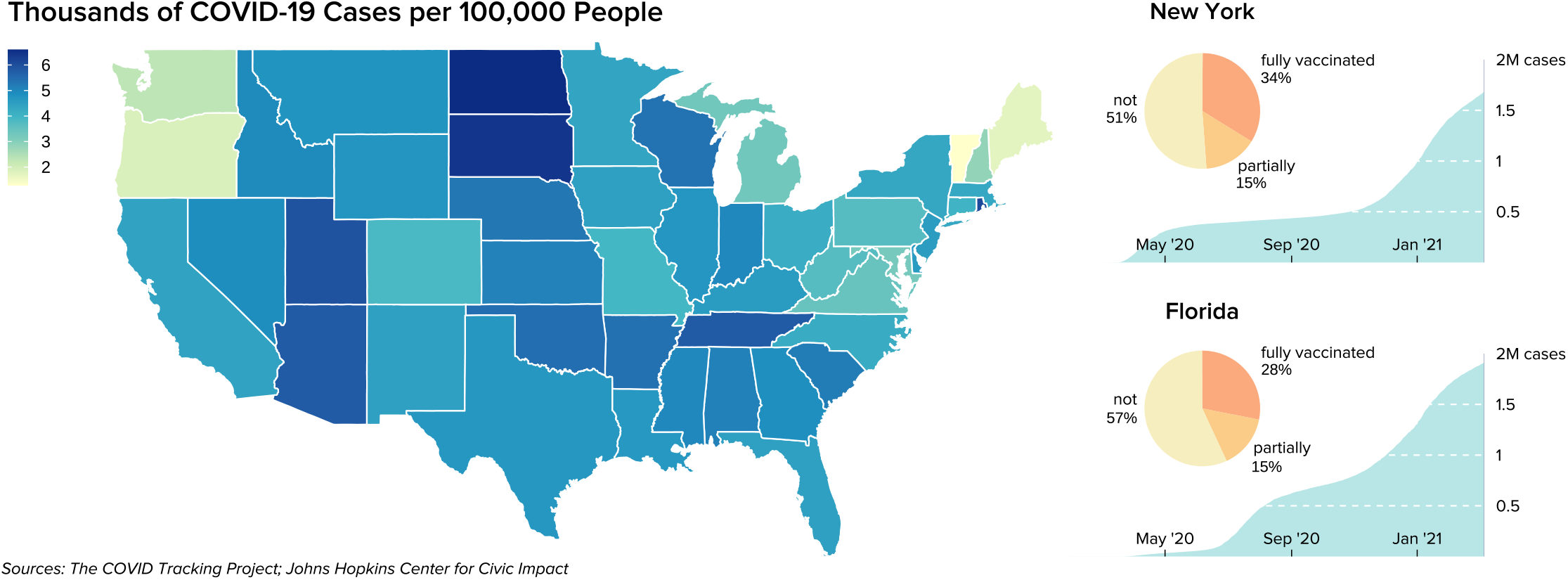
Precise arrangement of ggplot2 objects with plotgardener. Five ggplot2 objects and additional text elements were arranged into this multi-panel figure using plotgardener. Left: A map of the United States depicts COVID-19 cases per 100,000 people in each state. Right: Pie charts depict state vaccination percentages and line plots describe cumulative COVID-19 cases in New York and Florida. The plotgardener functions used to create this figure include pageCreate, plotGG, and plotText. Code to reproduce this plot is included in the plotgardener package.

In summary, plotgardener provides a new paradigm for generating complex publication-quality figures of both genomic and non-genomic data types, making it an invaluable tool for R users and data scientists from virtually any discipline.

## Acknowledgments

We would like to thank Hyejung Won and Jason Stein for helpful discussions and feedback. We thank Muhammad Saad Shamim and Neva Durand for assistance with the strawr package. This work was supported by NIH grants (R35-GM128645 to D.H.P.). N.E.K. and E.S.D. were supported by the NIH-NIGMS training grant T32-GM067553. S.M.P. is supported by the NSF GRFP DGE-1650116. M.I.L. was supported by NIH grants R01-MH118349 and R01-HG009937.

## Author information

Nicole E Kramer: Conceptualization, Software, Methodology, Writing

Eric S Davis: Conceptualization, Software, Writing

Craig D Wenger: Software

Erika M Deoudes: Visualization

Sarah M Parker: Software

Michael I Love: Software, Writing

Douglas H Phanstiel: Conceptualization, Writing, Supervision, Funding acquisition

## Methods

### Visualization methods

Plotgardener is an open-source extension for R, building its visualization functions from primitive graphical functions in the grid package^30^. Each plot and annotation is drawn within its own defined graphical region, or viewport, and then placed on a larger plotgardener page. These viewports give the power to specify the size and placement of plot containers and clip data to precise genomic and data axis measurements. To obtain large, reference genomic annotation data, plotgardener integrates and utilizes packages and data objects through Bioconductor.

### Gene and transcript label publication frequency mining

Annotations for genes in PubMed articles were obtained from the PubTator text mining tool^31^ and counted for each unique gene ID. Publication frequencies were matched via gene ID to Bioconductor transcript database (TxDb) gene IDs for the 29 built-in plotgardener genomes.

### Evaluating runtimes of plotgardener plotting functions

To calculate plotgardener plotting runtimes, we used the R package microbenchmark^32^. plotHicSquare, plotSignal, plotGenes, and plotRanges functions were timed for various genomic region sizes and resolutions. Each condition was timed on 20 random genomic regions generated by BedtoolsR^33^.

## Data availability

Various publicly available datasets are included with a supplementary plotgardenerData package and were used to demonstrate the functionalities of plotgardener. Hi-C datasets from the GM12878 and IMR90 cell lines were downloaded from GEO^34^ under the accession code GSE63525. CTCF ChIP-seq signal files for the GM12878 and IMR90 cell lines were downloaded from the ENCODE portal^35^ with accession codes ENCFF312KXX and ENCFF603PYX. H3K27ac ChIP-seq signal files for the GM12878 and IMR90 cell lines were downloaded from the NIH Roadmap Epigenomics Project^36^ with reference epigenome identifiers E116 and E017. COVID-19 case data was downloaded from The COVID Tracking Project (https://covidtracking.com/). State population data and state COVID-19 vaccination data were downloaded from the Johns Hopkins Centers for Civic Impact COVID-19 GitHub repository (https://github.com/govex/COVID-19/).

## Notes

https://phanstiellab.github.io/plotgardener/

https://github.com/PhanstielLab/plotgardener

### Competing Interest Statement

The authors have declared no competing interest.

